# Rho of plant GTPase MxROP1 regulates Fe deficiency responses by targeting Zinc Ribbon 3 in apple rootstock

**DOI:** 10.1101/2022.07.25.501490

**Authors:** Keting Li, Longmei Zhai, Lizhong Jiang, Qiran Sun, Ting Wu, Xinzhong Zhang, Xuefeng Xu, Zhenhai Han, Yi Wang

## Abstract

Small G protein is a multifunctional molecular switch that can regulate plant growth, development and responses to the environment. However, how Rho-related GTPase of plants (ROPs) regulates the response to Fe deficiency has not been well clarified. Here, we found that Fe deficiency induced MxROP1 in *Malus xiaojinensis* at both the transcriptional and translational levels. The overexpression of MxROP1, MxROP1^DN^ (inactive form) and MxROP1^CA^ (active form) in apple roots increased the activity of ferric chelate reductase and the ability to acidify the rhizosphere, and lines that overexpressed MxROP1^DN^ exhibited the strongest reaction to enhance Fe uptake. Yeast two-hybrid library screening indicated that MxROP1 interacted with ZR3.1, a DNL zinc finger protein that negatively regulates Fe deficiency responses. We further identified their interaction *in vitro* and *in vivo* using pull-down and bimolecular fluorescence complementation assays, respectively, and MxROP1^DN^-MxZR3.1 interacted the most strongly. Furthermore, MxROP1 negatively affected the stability of MxZR3.1 protein *in vitro* as shown by a cell semi-degradation assay, and the application of MG132 inhibited the degradation of MxZR3.1-HIS proteins. This indicated that MxROP1 caused the degradation of MxZR3.1 protein through the 26S proteasome pathway. Similar results were found in OE-MxROP1+OE-MxZR3.1 transgenic apple callus compared with those in the OE-MxZR3.1 callus. We also demonstrated that MxZR3.1 interacted with MxbHLH39, a known positive transcription factor and core component of Fe deficiency, and MxROP1 affected the interaction of MxZR3.1-MxbHLH39 using a competitive binding assay. This illuminated one MxROP1-MxZR3.1-MxbHLH39 pathway that maintains Fe homeostasis in *M. xiaojinensis*.

## INTRODUCTION

Apple (*Malus domestica*) is an important horticultural and economically valuable fruit tree. In apple production, the desired cultivars are grafted onto the rootstock, which is responsible for absorbing nutrients. Various nutrients affect growth and development of apple trees, including macro- and micro-elements. Iron (Fe) is an indispensable micronutrient and also one of the most important factors in plants. Although Fe is abundant in the earth’s crust, soluble iron that can be absorbed by plants is highly limited. Fe deficiency affects plant productivity and crop yields (Briat et al., 2015). Many plants, including apple, citrus and some vegetables, suffer from Fe deficiency chlorosis in neutral or alkaline soils (Kobayashi and Nishizawa, 2012; Lei et al., 2020).

To adapt to an environment deficient in iron, plants have evolved two mechanisms (Römheld and Marschner, 1986). Strategy I involves reduction and is primarily utilized by dicotyledons and non-gramineous monocotyledons. Strategy II involves chelation and primarily occurs in monocotyledons (Marschner et al., 1986). When the plants that utilize strategy I grow in calcareous or arid soil, they usually activate a series of functional proteins, such as PLASMA MEMBRANE PROTON ATPASE 2 (AHA2), FERRIC REDUCTASE OXIDASE2 (FRO2) and IRON TRANSPORTER1 (IRT1), which improve the dissolution, reduction and absorption of environmental Fe, respectively (Römheld and Marschner, 1986; Robinson et al., 1999). This process is strictly regulated by many transcription factors at the transcriptional level, such as FER-LIKE IRON DEFICIENCY-INDUCED TRANSCRIPTION FACTOR (FIT) (Colangelo and Guerinot, 2004). FIT is a bHLH transcription factor that interacts with one of the four members of the bHLH Ib subgroup (bHLH38, bHLH39, bHLH100 and bHLH101) to regulate the response to Fe deficiency (Yuan et al., 2008; Wang et al., 2013). Normally, bHLH transcription factors form heterodimers with other regulatory proteins to affect the genes related to Fe absorption under Fe deficiency, including *IRT, FIT*, and *HAs* (Vert et al., 2002; Colangelo and Guerinot, 2004; Ivanov et al., 2012). Some other transcriptional factors have also been identified in apple, including *MdbHLH104* (the homologous gene of *AtbHLH104*), *MxMYB1* (the homologous gene of *AtMYB1*) and *MxERF4* (the homologous gene of *AtERF4*) (Zhao et al., 2016; Shen et al., 2008; Zhang et al., 2020).

Zinc finger protein ZFPs are types of transcription factors that are involved in plant growth and development and resistance to biotic and abiotic stresses (Le et al., 2016). Under Fe deficiency stress, one ZFP member ZAT12 in *Arabidopsis thaliana* roots is involved in the regulation of stability of the FIT protein under the hydrogen peroxide (H_2_O_2_) pathway (Le et al., 2016). Previous studies have found a class of C4 zinc finger protein ZRs in apple rootstocks, but the biological function of these proteins in response to Fe deficiency in apple has not been well characterized. Moreover, a class of short C-terminal motif (DNL) zinc finger proteins with a D (D/H) L motif are localized in plastids and mitochondria in higher plants. They are members of a systematically conserved small zinc ribbon (ZR) protein family. The ZR protein was first identified as a DNA-binding motif in the TFIIIA transcription factor from African clawed frog (*Xenopus laevis*) (Miller et al., 1985; Lee et al., 1989). These proteins have only been reported in plants, but their potential functions still remain unclear.

Plant small G protein ROPs are a class of highly conserved molecular switches that are ubiquitous in eukaryotes and participate in a variety of signaling pathways, including cell growth, development and responses to the environment (Limor et al., 2003; Xu et al., 2010; Xu et al., 2014; Zheng et al., 2002). There are 11 types of ROPs in *A. thaliana* (Berken and Wittinghofer, 2008). Among them, AtROP6 has been identified to positively regulate Fe deficiency responses (Zhai et al., 2018). In other species, research on the function of ROP primarily focuses on biological stress. For example, silencing MtROP9 affects the interaction between pathogenicity and symbiosis (Kiirika et al., 2012). MdROP8 in apple is involved in SRNase-mediated self-incompatibility (Meng et al., 2014). ROPs rely on the cycle between their own active (CA) and inactive (DN) states to regulate downstream effectors and participate in many cellular activities (Berken and Wittinghofer, 2008). Overexpressing CA-LjROP6, a positive regulator of infection thread formation and nodulation, promoted root hair curling and increased the infection events and nodule number, while DN-LjROP6-ox exhibited an opposite phenotype (Ke and Peng, 2019). In addition, the overexpression of EgROPl, EgROP1-CA and EgROP1-DN altered the cellular morphology in leaves, but the alternating rectangular and spindle-shaped cells and smaller circular or triangular-shaped cells of EgROP1-DN-OX appeared different from those of EgROP1-CA and EgROP1-DN (Camille et al., 2009). The CA and DN states of ROP may play similar or opposite roles in some specific physiological processes, but whether their different states function during Fe uptake merits further study.

In plants, ROP-guanosine triphosphate (GTP) transmits signals downstream through protein-protein interactions, and most ROP effectors play a direct role in cell tissues (Han et al., 2018). We previously reported that MxROP1 from the iron efficient apple rootstock *M. xiaojinensis* is involved in Fe deficiency responses (Zhai et al., 2021), but how it regulates Fe uptake merits further study. Here, we showed that the overexpression of *MxROP1, MxROP1*^*CA*^, and *MxROP1*^*DN*^ improved the absorption of Fe in apple rootstock under normal Fe conditions. By screening a yeast library and performing bioinformatics, one protein zinc ribbon 3 (ZR3.1) was identified that interacted with MxROP1^DN^. MxZR3.1 is a typical zinc finger protein (Pfam: zf-dnl) with a C-terminal C4-type ZR domain, which negatively regulates Fe uptake by binding MxbHLH39 in apple rootstock, while the involvement of MxROP1 reduced the negative regulatory function of MxZR3.1. In summary, we provide a new mechanism about the participation of MxROP1 and MxZR3.1 to maintain iron homeostasis in apple rootstock.

## RESULTS

### Characterization and subcellular localization of MxROP1

The Fe-sufficient genotype apple rootstock *M. xiaojinensis* was subjected to Fe deficiency treatment. Compared with plants under Fe sufficiency, Fe deficiency treatment significantly induced the activity of ferric chelate reductase (FCR) in roots (Figure 1**A** and **B**). The rhizosphere pH was obviously lower, and H^+^-ATPase activity increased significantly under Fe deficiency conditions (Figure 1**C** and **D**). The small G protein ROPs have been reported to play important roles in the adaptive responses to Fe deficiency (Zhai et al., 2018; Zhai et al., 2021). We finally identified 10 ROPs in the apple genome, and the lengths of their amino acid varied from 21 to 23 kDa (Table S1). The expression of *MdROPs* in different tissues was investigated using public RNA-Seq data (http://bioinformatics.cau.edu.cn/AppleMDO/index.php) (Da et al., 2019). We found that the levels of expression of *MdROP2, 3, 7, 8*, and *10* were higher than those of the other members in flower, stem and mature leaf tissues, while only *MdROP5* was expressed at its lowest level in young leaves (Figure S1**A**). We then analyzed their expression in *M. xiaojinensis* roots under Fe deficiency treatment, and *MdROP3, 5, 6, 7*, and *8* were dramatically induced (Figure S1**B**). A phylogenetic analysis indicates that *MdROP7* clusters with *AtROP6* (Figure S1**C**), which has been demonstrated to positively regulate Fe deficiency responses in *A. thaliana* (Zhai et al. 2018). Moreover, this *MdROP7* is the gene that has been identified as *MxROP1* in *M. xiaojinensis* (Figure S1**D**) (Zhai et al., 2021). Therefore, *MxROP1* (*MdROP7*) was used for further research in this study.

**Figure 1.**
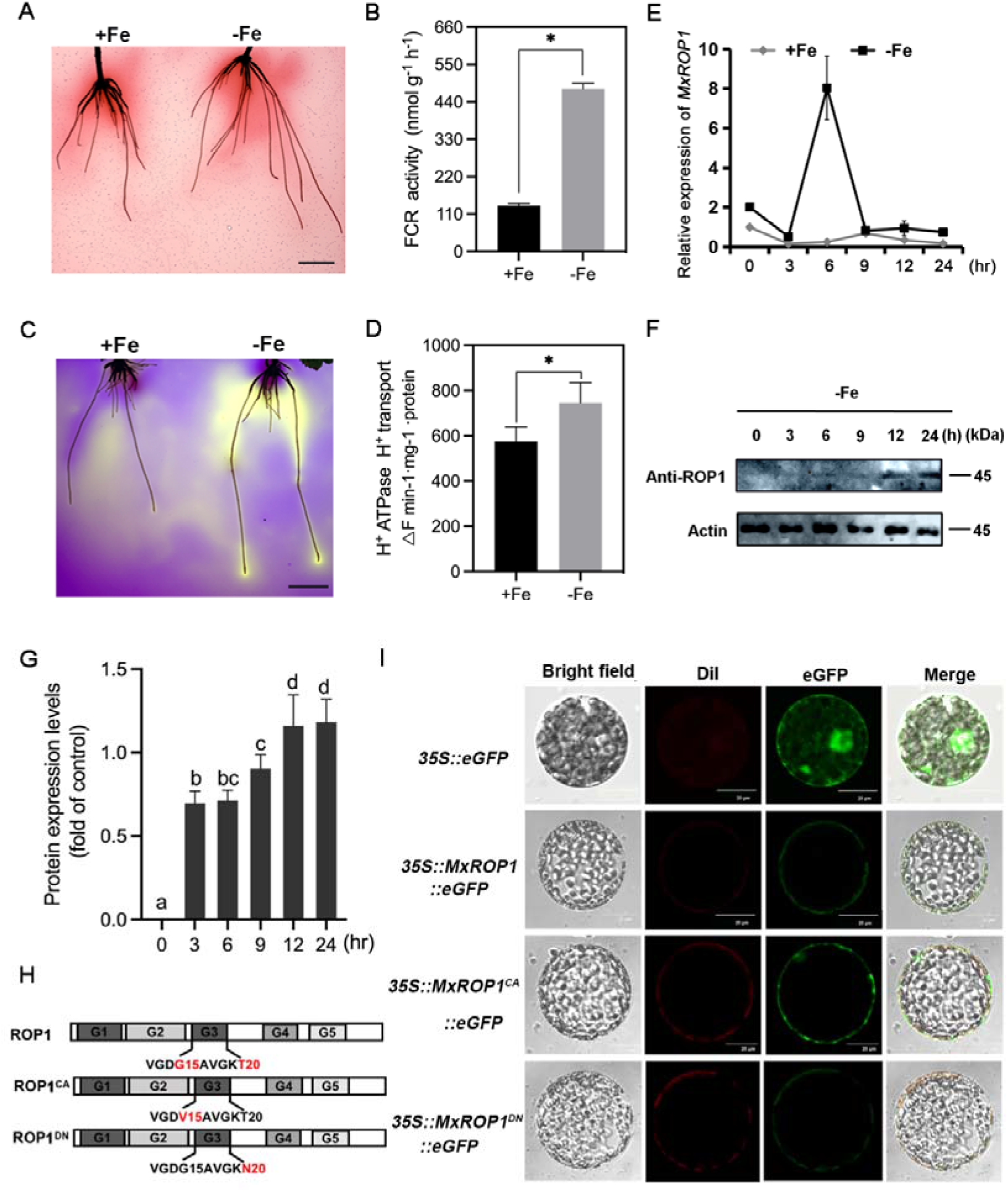
Identification of MxROP1 as a candidate factor in response to Fe deficiency in *Malus xiaojinensis*. **A-B**. FCR activity in the roots of *Malus xiaojinensis* increased significantly under Fe-deficient conditions for 5 days as shown by qualitative detection (A) and quantitative detection (B). Data are shown as the means ± SD (n=3). Bar=1 cm. **C**. The acidification of rhizosphere in *M. xiaojinensis* was enhanced under Fe deficiency when compared with that under normal Fe conditions. Bar=1 cm. **D**. H^+^-ATPase activity was induced in the roots of *M. xiaojinensis* under Fe deficiency. **E**. Expression of *MxROP1* in the roots of *M. xiaojinensis* was dramatically induced at 6 h after Fe deficiency treatment, while there was no change under Fe normal conditions. Data are shown as the means ± SD (n=3). **F**. Levels of MxROP1 protein were induced in the roots of *M. xiaojinensis* under Fe deficiency using MxROP1 specific antibodies. ACTIN was used as a loading control. **G**. The level of MxROP1 protein significantly increased in the roots of *M. xiaojinensis* under Fe deficiency. The experiments were conducted in triplicate, and the value relative to 0 h of protein level was established as “1.” Data are shown as the means ± SD. **H**. Schematic diagrams of MxROP1 and its mutational site. **I**. MxROP1, MxROP1^DN^, and MxROP1^CA^ were localized on the cell membrane in *Nicotiana benthamiana* leaf protoplasts. Bar=20 μm. *a significant difference between the +Fe and -Fe conditions was determined using a two-tailed Student’s *t*-test with pooled variance. Bars with different letters are significantly different at P < 0.05 (ANOVA, Duncan correction). The bars show standard deviations. ANOVA, analysis of variance; FCR, ferric chelate reductase; SD, standard deviation.

The level of expression of *MxROP1* in *M. xiaojinensis* roots was detected at different treatment times of Fe deficiency. It showed no significant differences under Fe normal conditions but was significantly induced by approximately 4-fold at 6 h of Fe deficiency conditions (Figure 1**E**). In addition, we customized a specific antibody of MxROP1 to verify whether the MxROP1 protein was affected by Fe deficiency. The MxROP1 protein gradually accumulated with the extension of time of Fe deficiency (Figure 1**F** and **G**). These results showed that the roots of *M. xiaojinensis* MxROP1 responded to Fe deficiency at both the transcriptional and translational levels. Small GTPases play roles by switching between the active state (GTP-bound) and the inactive form that binds GDP (Rivero et al, 2019). We obtained constitutively active GTP-bound ROP1 (MxROP1^CA^) and the dominant negative nucleotide-free ROP1 (MxROP1^DN^) using site-directed mutagenesis (Molendijk et al, 2001) (Figure 1**H**). A subcellular localization analysis indicated that all three proteins were located on the cell membrane (Figure 1**I**).

### Physiological responses of the overexpression lines of *MxROP1, MxROP1*^*CA*^ and *MxROP1*^*DN*^ to Fe deficiency

To investigate the potential mechanism of the regulation of MxROP1 in Fe absorption, we obtained several overexpression lines of *MxROP1, MxROP1*^*CA*^ and *MxROP1*^*DN*^ using *Agrobacterium rhizogenes*-mediated transformation technology (Figures S2 and S3). Under Fe normal conditions, the activity of FCR in the roots of the three overexpressed lines increased significantly, and MxROP1^DN^ responded the most strongly (Figure 2**A** and **B**). Similarly, rhizosphere acidification and H^+^-ATPase activity were both induced in the three transgenic lines, particularly in the MxROP1^DN^ overexpressed lines, when compared with that in control plants under normal Fe conditions (Figure 2**C** and **D**). Furthermore, Perls staining showed that the transgenic roots accumulated more iron compared with the control (Figure 2**E**), indicating that both the active and inactive forms of MxROP1 positively regulate Fe absorption.

**Figure 2.**
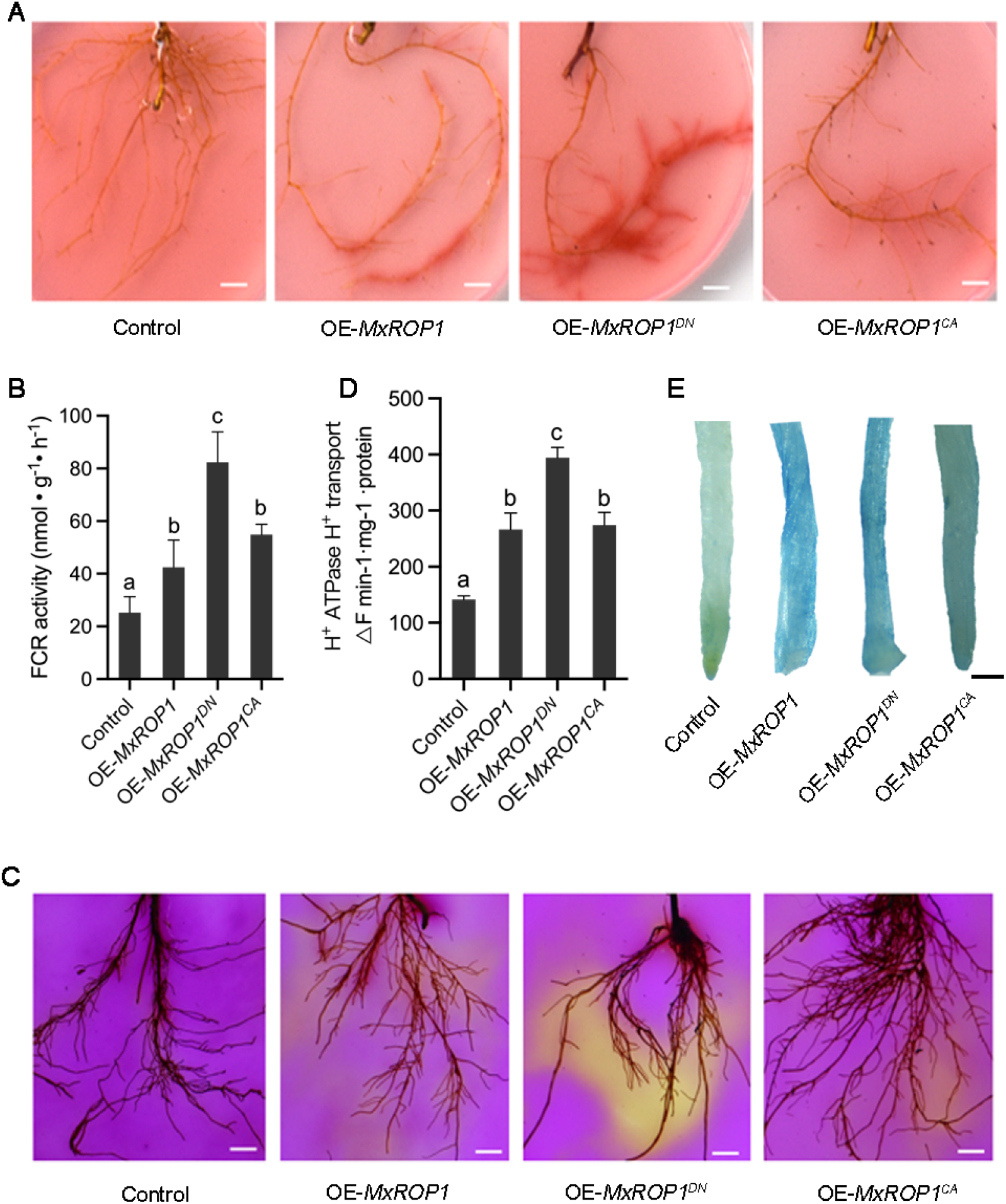
Overexpression of MxROP1, MxROP1^DN^, and MxROP1^CA^ in apple plants with enhanced Fe deficiency responses. **A-B**. FCR activity in the roots of OE-MxROP1, OE-MxROP1^DN^, and OE-MxROP1^CA^ plants was significantly enhanced when compared with that in control plants under Fe normal conditions. Bar=0.5 cm. Data are shown as the means ± SD (n=3). **C-D**. Rhizosphere acidification in the roots of OE-MxROP1, OE-MxROP1^DN^, and OE-MxROP1^CA^ plants increased dramatically under Fe normal conditions. The qualitative detection of acidification is indicated by a yellow color around the roots. Data are shown as the means ± SD (n=3). Bar=1 cm. **E**. Fe content in the roots of MxROP1-OE, MxROP1^DN^-OE, and MxROP1^CA^-OE visibly increased as indicated by Perls staining under Fe normal conditions. Bar=200 µm. The bars show standard deviations. ANOVA, analysis of variation; FCR, ferric chelate reductase; SD, standard deviation.

### MxROP1 associates with MxZR3.1 via direct binding

To investigate the MxROP1-mediated pathway in response to Fe absorption, MxROP1 was used as a protein-bait to screen the potential proteins that interacts with using the Fe deficiency cDNA yeast two-hybrid (Y2H) library of *M. xiaojinensis*. A DNL zinc finger protein ZR3.1 (GenBank accession number: MD01G1105600) that contained the classical zinc finger domain zf-DNL and EAR domain was screened to interact with MxROP1. A multi-sequence alignment and phylogenetic analysis then showed that MxZR3.1 clustered to AtZR3 and MdZR3.1 (Figure S5**A** and **B**). An additional Y2H assay further demonstrated that MxZR3.1 and MxROP1 interact *in vitro* (Figure S4).

To further identify the association of MxROP1 with MxZR3.1, a DUAL membrane yeast two-hybrid assay was performed. We found that MxROP1 and MxROP1^DN^ were able to interact with MxZR3.1, while MxROP1^CA^ could not interact with MxZR3.1 (Figure 3**A**). We used the Y2H assays for additional experiments to determine whether other MxZRs could interact with MxROP1, but there was no interaction (Figure S5**C**). In addition, MxZR3.1 associated with MdROP3, 6 and 7 (Figure S5**D**), which clustered together, indicating that MxZR3.1 could interact with ROP members of the same branch (Schepetilnikov et al., 2017). In contrast, a pull-down assay showed that both GTP- and GDP-bound form of MxROP1 interacted with MxZR3.1 *in vivo* (Figure 3**B**). The bimolecular fluorescence complementation (BiFC) assay showed similar results (Figure 3**C**). However, it indicated that the interaction between MxROP1^CA^ and MxZR3.1 was weak (Figure 3**B** and **C**), indicating that the GDP-bound form of MxROP1 (MxROP1^DN^) strongly interacted with MxZR3.1.

**Figure 3.**
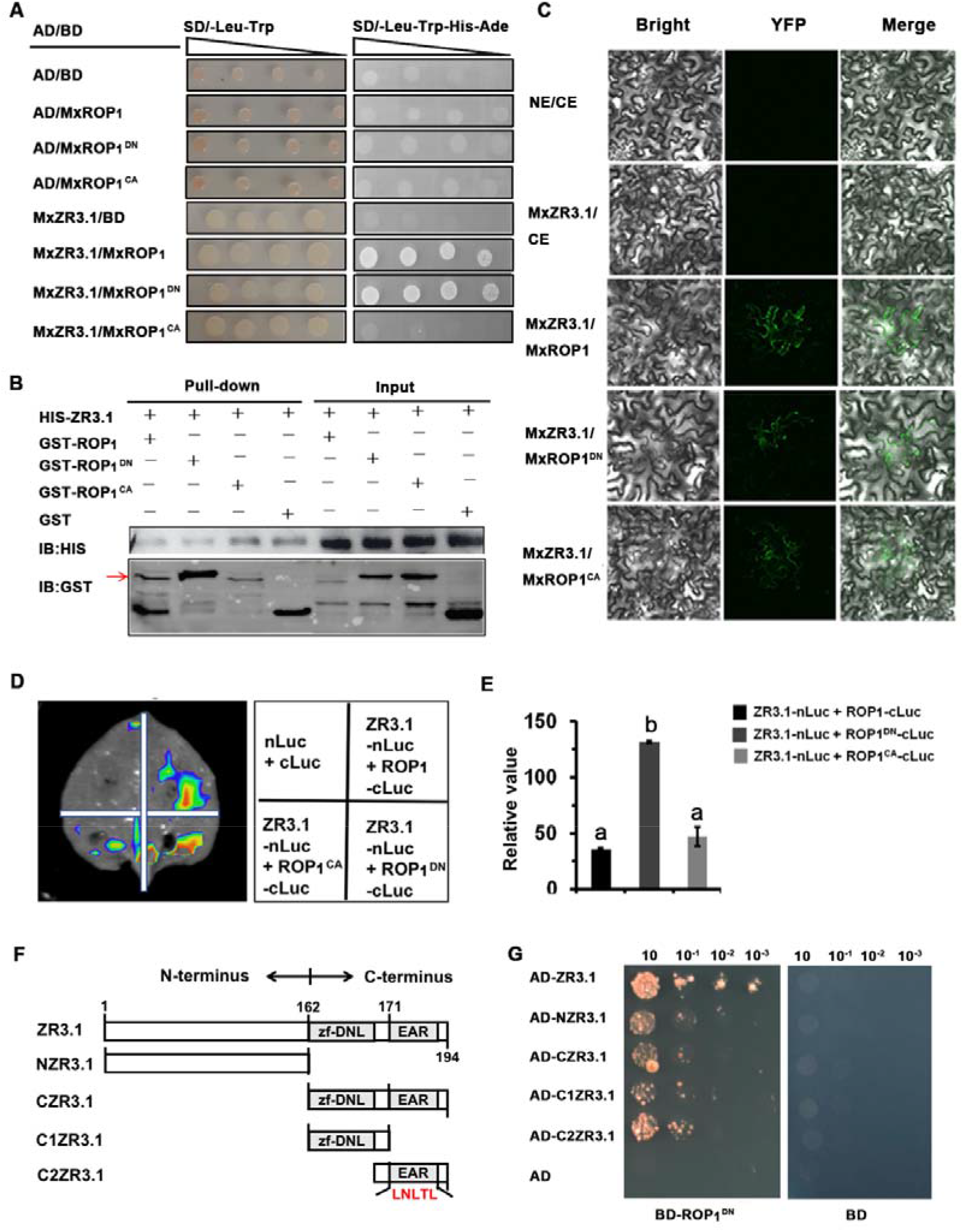
MxROP1, MxROP1^DN^, and MxROP1^CA^ interact with the MxZR3.1 protein. **A**. MxROP1 and MxROP1^DN^ interacted with MxZR3.1, but MxROP1^CA^ did not interact in the Y2H assay. **B**. The pull-down assay showed that the ROP1-, ROP1^DN^-, and ROP1^CA^-GST proteins interacted with the ZR3.1-HIS protein. The red arrow indicates the recombinant ZR3.1-His protein. **C**. MxROP1, MxROP1^DN^, MxROP1^CA^ interacted with MxZR3.1 in tobacco (*Nicotiana benthamiana*) leaves using a BiFC assay. Bar=200 μm. **D**. MxROP1, MxROP1^DN^, and MxROP1^CA^ interact with MxZR3.1 in *N. benthamiana* leaves as shown by an LCI assay. The 35S:MxZR3.1-nLUC and 35S::cLUC-MxROP1, 35S::MxROP1^DN^, and 35S::MxROP1^CA^ constructs were co-expressed in tobacco leaves and then imaged with the LUC signal after 72 h of transfection. **E**. Quantification of the fluorescence intensity in the leaves of the LCI assay showed that the signal of MxZR3.1 and MxROP1^DN^ coexpression was the strongest. Data are shown as the means ± SD (n=3). **F**. Schematic representation of the functional domains of MxZR3.1 (N terminal and C terminal basic zf-DNL. The EAR domains are indicated). **G**. Y2H: ZR3.1 and its N-terminal domain (NZR3.1), C-terminal (CZR3.1), zf-DNL domain (C1ZR3.1) and EAR domain (C2ZR3.1) interacted with ROP1^DN^. AD-ZR3.1, -NZR3.1, -CZR3.1, -C1ZR3.1 and -C2ZR3.1 were assayed for interaction with BD-ROP1^DN^ as indicated. Bars with different letters are significantly different at P < 0.05 (ANOVA, Duncan correction). The bars show standard deviations. ANOVA, analysis of variance; BiFC, bimolecular fluorescence complementation; GST, glutathione-S-transferase; LCI, luciferase complementation imaging; LUC, luciferase; Y2H, yeast two-hybrid; YFP, yellow fluorescent protein.

Moreover, luciferase complementation imaging (LCI) assays showed that the interaction between ZR3.1 and MxROP1^DN^ was stronger than that with MxROP1 or MxROP1^CA^ (Figure 3**D** and **E**). The strong interaction of MxROP1^DN^ with MxZR3.1 led us to seek the exact binding site. Next, we analyzed the conserved domains of MxZR3.1 and obtained four fragments that included the N-terminal half of MxZR3.1 (NZR3.1) and the C-terminal half of MxZR3.1 (CZR3.1) and C1ZR3.1 and C2ZR3.1 (Figure 3**F**). Surprisingly, the Y2H system indicated that MxROP1^DN^ is bound to NZR3.1, CZR3.1, C1ZR3.1 and C2ZR3.1 (Figure 3**G**). These results implied that the inactive MxROP1 played a more important role in interacting with MxZR3.1.

### MxZR3.1 negatively regulates Fe deficiency responses

We observed that the transcriptional level of *MxZR3*.*1* was induced by Fe deficiency (Figure S6**A**), while its protein level was reduced within 24 h under Fe deficiency (Figure S6**B**). Therefore, we hypothesized that MxZR3.1 may respond to Fe deficiency in *M. xiaojinensis*. Since there is a classical transcriptional inhibitory domain EAR in the *MxZR3*.*1* sequence, we hypothesized that *MxZR3*.*1* may play an inhibitory role in the Fe deficiency response pathway.

Next, we silenced *MxZR3*.*1* in the apple rootstocks *M. baccata* using tobacco tobacco rattle virus (TRV) silencing technology. The expression of *MxZR3*.*1* decreased in the roots of TRV-*MxZR3*.*1* plants (Figure 4**A**). The activity of FCR in the roots of TRV-*MxZR3*.*1* increased significantly under Fe deficiency (Figure 4**B**). Similarly, the rhizosphere of TRV-*ZR3*.*1* plants was obviously acidified, particularly without iron, when compared with that in TRV plants, indicating that MxZR3.1 affected the release of protons induced by Fe deficiency (Figure 4**C**). Moreover, silencing MxZR3.1 increased the content of iron in roots of apple rootstocks (Figure 4**D**). At the molecular level, we detected the expression of Fe-absorption genes *IRT1* and *FRO2*, which were clearly enhanced in the roots of TRV-*ZR3*.*1* plants under Fe deficiency compared with those in the TRV plants (Figure 4**E**-**F**). In addition, *FIT* and *ZAT12c*, additional major transcription factors that respond to Fe deficiency, were highly expressed in TRV-*ZR3*.*1* under Fe-sufficient conditions (Figure S7**A** and **B**), illustrating that MxZR3.1 negatively modulated Fe deficiency responses.

**Figure 4.**
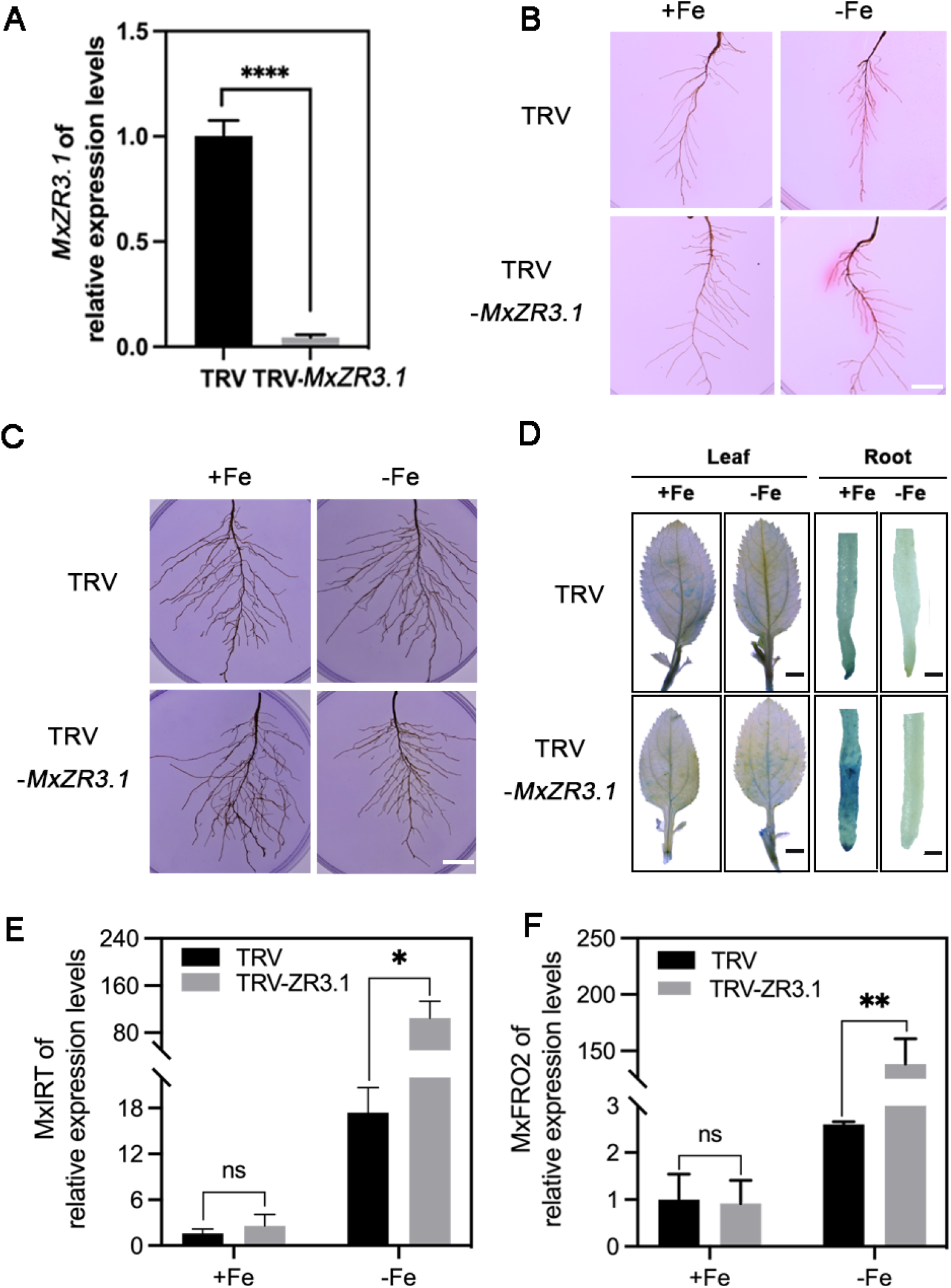
Silencing *MxZR3*.*1* in apple rootstocks increased the Fe deficiency response. **A**. The expression of *MxZR3*.*1* was significantly reduced in *MxZR3*.*1-*silenced *Malus baccata* (L) Borkh. apple rootstock (TRV-*MxZR3*.*1*). The level of expression in the TRV line was established as “1,” and the data are shown as means ± SD (n=3). **B**. FCR activity in the roots of TRV-*ZR3*.*1* lines obviously under Fe deficiency conditions when compared with that in the TRV lines. Bar=0.5 **C**. Rhizosphere acidification of TRV-*ZR3*.*1* lines was clearly enhanced when exposed to +Fe or -Fe for 3 d compared with that of the TRV lines. Acidification was indicated by a yellow color around the roots. Bar=0.5 cm. **D**. Perls staining showed that the Fe content in roots of TRV-*MxZR3*.*1* under +Fe conditions was higher than that in the of TRV plants. Bar=1 cm (Leaf); Bar=200 µm (Root). **E-F**. The levels of transcription of Fe response genes *MxIRT1* (E) and *MxFRO2* (F) in the TRV and TRV-*ZR3*.*1* lines under +Fe or -Fe. The level of expression in the TRV line was established as “1,” and the data are shown as the means ± SD (n=3). *a significant difference between the TRV and TRV-MxZR3.1 lines, as determined using a two-tailed Student’s *t*-test with pooled variance. FCR, ferric chelate reductase; SD, standard deviation; TRV, tobacco rattle virus.

### Overexpressing MxROP1 mitigates the negative iron deficiency responses regulated by MxZR3.1

The fact that *OE-MxROP1* and TRV-*MxZR3*.*1* lines enhanced the Fe deficiency responses showed that *MxROP1* and *MxZR3*.*1* played opposite roles in the absorption of Fe by *M. xiaojinensis*, but how their interaction regulated Fe deficiency responses merits further identification. Therefore, we obtained transgenic apple callus, including *OE-MxZR3*.*1* and *OE-MxZR3*.*1*+*OE-MxROP1* (*MxROP1* overexpressed in *OE-MxZR3*.*1* apple callus), using *A. tumefaciens* mediated transformation (Figure 5**A** and **B**).

**Figure 5.**
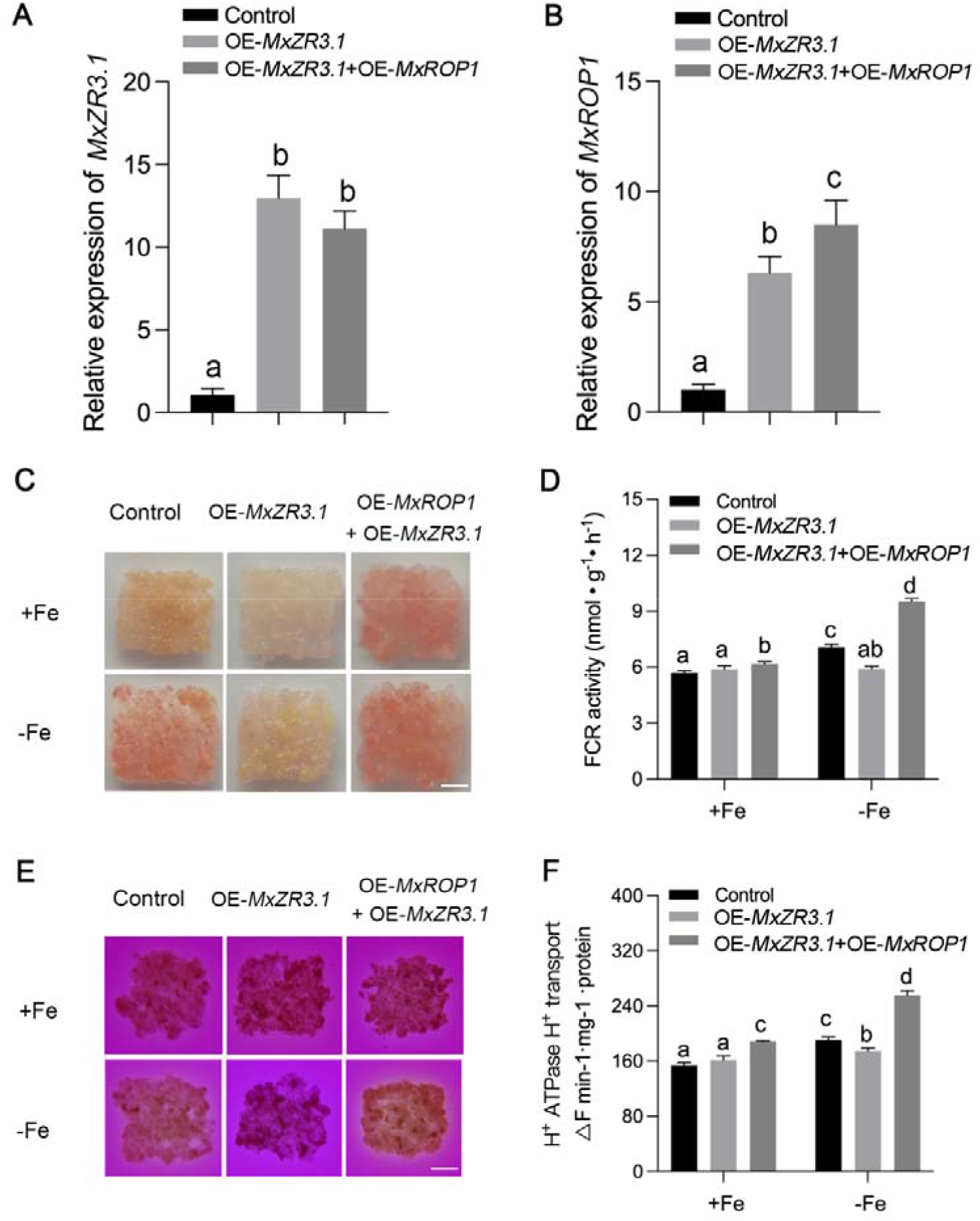
MxROP1 binds to MxZR3.1 to enhance the Fe deficiency responses. **A-B**. The expression of *MxZR3*.*1* (A) and *MxROP1* (B) in control (wild type) callus, *OE-MxZR3*.*1* transgenic callus and *OE-MxZR3*.*1*+*OE-ROP1* co-transgenic callus. **C**. Visualization of FCR activity in OE-MxZR3.1 obviously decreased, while the overexpression of MxROP1 in OE-MxZR3.1 callus inhibited the decreased FCR activity under Fe deficiency conditions. **D**. FCR activity in the callus of OE-MxROP1+OE-MxZR3.1 was significantly enhanced compared with that in the control and OE-MxZR3.1 under Fe deficiency. Data are expressed as the mean ± SD (n=3). **E**. Rhizosphere acidification in the *OE-MxZR3*.*1* callus clearly decreased, while it was enhanced in the *OE-MxZR3*.*1*+*OE-MxROP1* co-transgenic callus. Acidification is indicated by a yellow color around the callus. **F**. Rhizosphere acidification in the *OE-MxZR3*.*1*+*OE-MxROP1* co-transgenic callus was strongly induced under +Fe and -Fe conditions compared with that in the control and *OE-MxZR3*.*1* callus. Data are expressed as the mean ± SD (n=3). Bars with different letters are significantly different at P < 0.05 (ANOVA, Duncan correction). The bars show standard deviations. ANOVA, analysis of variance; FCR, ferric chelate reductase; SD, standard deviation.

Compared with the control callus (wild type [WT]), FCR activity in the *OE-MxZR3*.*1* transgenic callus decreased notably under Fe deficiency (Figure 5**C** and **D**), and its rhizosphere acid induced a yellow color that clearly showed inhibition (Figure 5**E** and **F**). FCR activity and the rhizosphere acidification capacity were enhanced when MxROP1 was overexpressed in the OE-MxZR3.1 transgenic apple callus under Fe deficiency (Figure 5**C**-**F**). Moreover, the levels of expression of *MdIRT1, MdFRO2*, and *MdFIT* increased in co-transformed callus and decreased in *OE-MxZR3*.*1* callus (Figure S8**A-C**). These results demonstrate that MxROP1 interacts with MxZR3.1 to release the inhibition of Fe deficiency responses mediated by MxZR3.1.

### MxROP1 decreases the protein stability of MxZR3.1

Based on the decreased protein level of MxZR3.1 in the roots of *M. xiaojinensis* under Fe deficiency (Figure S6**B**), we hypothesized that MxROP1 could affect the stability of MxZR3.1 to participate in Fe deficiency responses. To verify this hypothesis, we performed a cell semi-degradation assay using the recombinant protein Myc-MxZR3.1 and the total protein of transgenic callus 35S::GFP and *35S::MxROP1*-GFP with or without the 26S proteasome complex inhibitor MG132. This showed that the Myc-MxZR3.1 protein degraded more quickly in the *35S::MxROP1*-GFP callus compared with the 35S::GFP callus (Figure 6**A**). In addition, treatment with MG132 significantly inhibited the degradation of Myc-MxZR3.1 in both 35S::GFP and *35S::MxROP1*-GFP (Figure 6**B**), suggesting that MxROP1 decreased the stability of MxZR3.1 through an ubiquitin-proteasome system *in vitro*.

**Figure 6.**
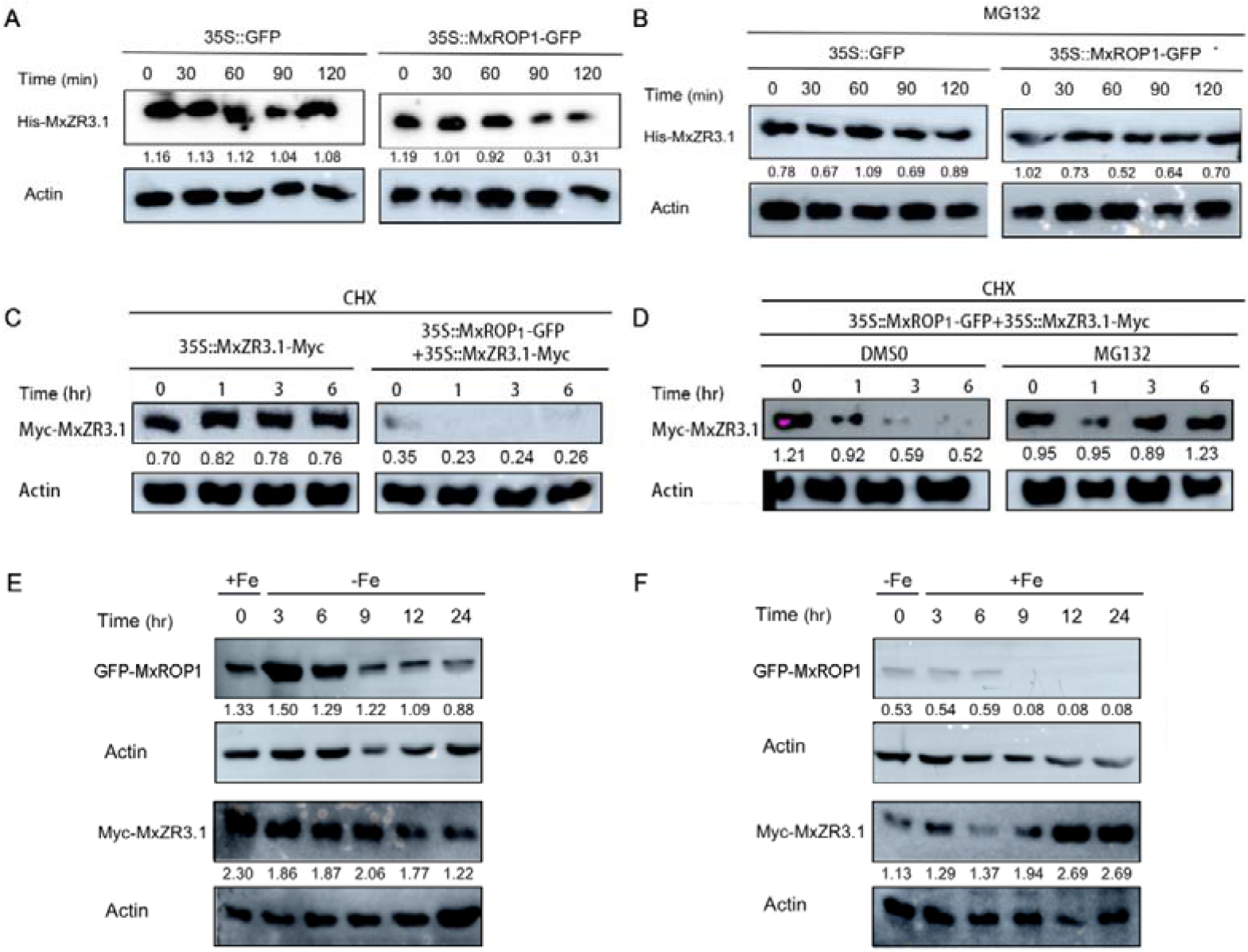
MxROP1 negatively affects the stability of MxZR3.1 protein. **A-B**. The purified recombinant His-MxZR3.1 protein degraded more quickly in the protein extract of 35S::MxROP1-GFP transgenic apple callus than in the 35S::GFP callus as shown by the cell-free degradation assays (A). Treatment with the proteasome inhibitor MG132 inhibited the quick degradation of His-MxZR3.1 protein in 35S::MxROP1-GFP transgenic apple callus (B). The levels of His-MxZR3.1 were visualized by immunoblotting using the anti-His antibody. ACTIN was used as the loading control. **C-D**. Under CHX treatment, the level of MxZR3.1-Myc protein decreased significantly in the *35S::MxZR3*.*1-Myc*+*35S::MxROP1* transgenic callus compared with that in the *35S::MxZR3*.*1-Myc* transgenic apple callus (C), while the application of MG132 inhibited MxZR3.1-Myc protein destabilization in the *35S::MxZR3*.*1-Myc*+*35S::MxROP1* transgenic callus (D). The total proteins were extracted at the times indicated and then analyzed by immunoblotting using the anti-Myc antibody. **E**. The level of MxROP1 protein increased, but the MxZR3.1 protein decreased at 3 h and 6 h after Fe deficiency, while both the MxROP1 and MxZR3.1 proteins decreased under prolonged Fe deficiency treatment. Transgenic apple callus were incubated with (+Fe) for 3 d, and then switched to no (-Fe+Frz) iron conditions. **F**. The level of MxROP1 protein gradually decreased, while the level of MxZR3.1 protein increased under Fe normal conditions. Transgenic apple callus were incubated without (-Fe+Frz) iron for 3 d and then switched to iron (+Fe) conditions. ACTIN was used as the loading control. CHX, chlorhexidine; DMSO, dimethyl sulfoxide.

We provided additional evidence that MxROP1 negatively regulated the accumulation of MxZR3.1 protein *in vivo*. The MxZR3.1-Myc protein degraded more quickly in the co-transformed callus (*35S::MxROP1-GFP* + *35S::MxZR3*.*1-Myc*) than in a single transformed callus (35S::MxZR3.1-Myc) after treatment with the translational inhibitor cycloheximide (CHX) (Figure 6**C**). Next, based on the CHX application, the co-transformed callus was treated with MG132 and dimethyl sulfoxide (DMSO) as a control. Treatment with MG132 significantly enhanced the accumulation of MxZR3.1-Myc protein in *35S::MxROP1-GFP* + *35S::MxZR3*.*1-Myc* callus, which was consistent with the results from cell semi-degradation assay (Figure 6**D**), indicating that the MxROP1-MxZR3.1 interaction resulted in the degradation of MxZR3.1-myc protein through the 26S proteasome pathway *in vivo*.

To detect the effects of MxROP1 on the abundance of MxZR3.1 protein in Fe absorption, *35S::MxROP1-GFP* + *35S::MxZR3*.*1-Myc* callus were subjected to Fe deficiency stress at different time periods. We found that the MxROP1 protein accumulated, while the abundance of MxZR3.1 protein was reduced at 3 h after Fe deficiency, but both MxROP1 and MxZR3.1 were degraded with the extension of Fe deficiency treatment, indicating that MxROP1 interacted with MxZR3.1 to affect its accumulation, thus, enhancing Fe absorption under Fe deficiency stress (Figure 6**E**. Surprisingly, the *35S::MxROP1-GFP* + *35S::MxZR3*.*1-Myc* callus were supplied with normal Fe after 3 days of Fe deficiency treatment. The level of MxROP1 protein decreased, but MxZR3.1 increased (Figure 6**F**), indicating that the MxROP1 protein may be degraded to release the MxZR3.1 protein to avoid excess Fe absorption under Fe normal conditions. These results demonstrated that MxROP1 was antagonistic to MxZR3.1 during the process of Fe homeostasis *in vivo*.

### MxROP1 competitively interacts with MxZR3.1 to release MxbHLH39

To explore how the ZR3.1 protein regulates Fe homeostasis in more detail, we conducted a Y2H assay and found that ZR3.1 interacted with MxbHLH39-1 (MD14G1086500) and MxbHLH39-2 (MD14G1086600), the homologous proteins of typical bHLH transcription factors in *Arabidopsis* that positively regulate the response to Fe deficiency (Yuan et al., 2008) (Figure 7**A** and **B**). We further analyzed the exact site of MxZR3.1 that interacted with MxbHLH39-1 and MxbHLH39-2, while only C1ZR3.1 weakly interacted with MxbHLH39-2 (Figure S9). In addition, a pull-down assay provided additional evidence that MxZR3.1 interacts with MxbHLH39-1 and MxbHLH39-2 *in vitro* (Figure 7**C**). BiFC assays showed that MxZR3.1 interacts with MxbHLH39-1 and MxbHLH39-2 in the nucleus (Figure 7**D**). These data demonstrate that MxZR3.1 interacts with MxbHLH39-1 and MxbHLH39-2.

**Figure 7.**
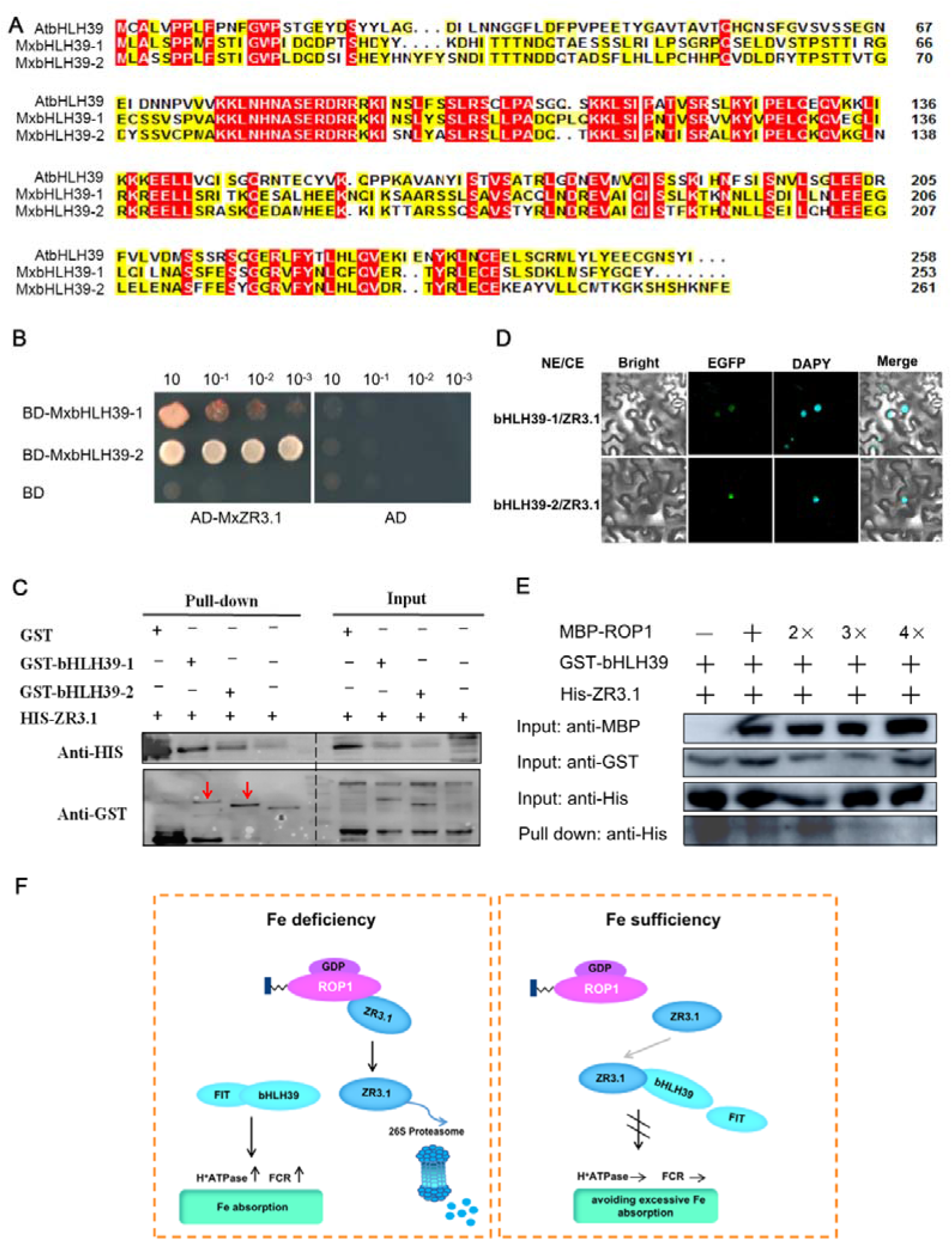
MxROP1 competes with MxbHLH39 to interact with MxZR3.1 to enhance the uptake of Fe. **A**. A multi-sequence alignment of MxbHLH39-1 protein with its orthologous protein MxbHLH39-2. **B**. A Y2H assay indicated that MxZR3.1 interacted with MxbHLH39-1 and MxbHLH39-2. **C**. A BiFC assay indicated that MxbHLH39-1 and MxbHLH39-2 interacted with ZR3.1 in *Nicotiana benthamiana* leaves. The co-transformation of cGFP-MxbHLH39-1, cGFP-MxbHLH39-2 and nGFP-ZR3.1 led to the reconstitution of GFP signal. DAPI was used to visualize the nuclei. The red arrow indicates the recombinant GST-bHLH39-1/2 protein. **D**. GST-tagged MxbHLH39-1 and MxbHLH39-2 interacted with HIS-tagged ZR3.1 as indicated on the left panel through the pull-down assay. **E**. Competitive binding assays showed that MxROP1 binds to MxZR3.1 to prevent the interaction of MxZR3.1 with MxbHLH39-1. A mixture of GST-MxbHLH39-1 and MBP-MxROP1 was added to immobilize His-MxZR3.1. The gradient indicates the increasing amount of MBP-MxROP1. **F**. A Proposed working model about the roles of MxROP1-MxZR3.1 in maintaining Fe homeostasis. BiFC, bimolecular fluorescence complementation; DAPI, 4’,6-diamidino-2-phenylindole; GFP, green fluorescent protein; Y2H, yeast 2-hybrid.

MxROP1 interacts with MxZR3.1 and acts oppositely in its responses to Fe deficiency, and bHLH39 is a positive transcription factor that regulates Fe absorption. This led us to estimate the effect of MxROP1 on the interaction between MxZR3.1 and MxbHLH39 using a competitive binding assay. First, we fused the MxbHLH39-1-GST protein induced *in vitro* to GST agarose, and MxROP1-MBP and MxZR3.1-HIS were then added. With the increase in MxROP1-MBP based on the concentration gradient, the interaction between bHLH39-1-GST and MxZR3.1-HIS gradually decreased (Figure 7**E**). These results indicate that MxROP1 is more competitive than MxbHLH39 in its interaction with MxZR3.1.

## DISCUSSION

The low solubility and bioavailability of iron in soil pose a serious threat to the biological growth of horticultural crops. Therefore, understanding the potential molecular mechanism of iron absorption and translocation will help to improve the efficiency of plants to utilize iron. It is well known that plants utilize complex sensing and signaling mechanisms to maintain iron homeostasis. Plant ROPs have been confirmed to be involved in Fe deficiency responses (Suzuki et al., 2011; Zhai et al., 2018), but only the ROS signaling pathway was reported to have a connection with the ROPs (Zhai et al., 2018; Zhai et al., 2021). In our study, to explore how the ROPs in apple rootstocks perceive Fe deficiency signals and transmit signals to enhance Fe uptake, we focused on the proteins that interact with ROPs. Here, we first reported that MxROP1 interacts with the Zinc finger protein MxZR3.1 to decrease its accumulation of protein, thus, releasing the MxbHLH39 protein to enhance Fe uptake under Fe deficiency in *M. xiaojinensis*. This finding provides significant insights into the mechanism of Fe homeostasis.

Plant small G protein ROPs play important roles in regulating numerous physiological processes (Yang et al., 2002; Carol et al., 2005). Normally, they function as protein molecular switches by alternately binding GTP or GDP (Mathur and Hü lskamp, 2002). In rice, CA-OsRac1 and DN-OsRac1 mutants activate and inhibit the induced defense of pathogens, respectively (Kawasaki et al., 1999). Moreover, the *ljrop* mutants and their CA or DN forms produced different root hair phenotypes (Liu et al., 2020). Here, overexpressing MxROP1, MxROP1^DN^ and MxROP1^CA^ enhanced the responses to iron deficiency. This is inconsistent with the previous results that found that CA and DN played opposite roles in pathogen defense (Kawasaki et al., 1999). However, treatment with DN-AtROP1 transformation lines in potato (*Solanum tuberosum*) resulted in plants that were more resistant to insects (Zhang et al., 2014), which is consistent with the finding that MxROP1^DN^ exhibited the strongest reaction to Fe deficiency. These differential responses could be owing to the differences between the mechanisms of different physiological processes. These differences could be explained by the concept that ROP GTPases act as molecular switches (Yang et al., 2002). ROPs in either the active or inactive state are strictly regulated to impact their function (Liu et al., 2020).

Two types of ROP effectors in plants have been reported, including ROP interactive CRIB motif-containing proteins (RICs) and interactor of constitutively active ROP1/ROP interactive partner 1 (ICR1/RIP1) (Wu et al., 2000; Lavy et al., 2006; Li et al., 2008). Here, we found a new zinc finger protein family member ZR3.1 that interacts with MxROP1. In addition to MdROP1 in the MdROPs family, MxZR3.1 can interact with MdROP3 and MdROP6, which are homologous to AtROP2 and AtROP4 in *A. thaliana*, respectively. AtROP2, AtROP4 and AtROP6 are members of the same subgroup, and some studies have confirmed that AtROP2, AtROP4 and AtROP6 interact with the same target protein (Schepetilnikov et al., 2017), which is consistent with the interaction of MxZR3.1 with MdROP3, MdROP6 and MdROP7 (MxROP1). However, MxROP1^DN^-MxZR3.1 interacted more strongly than MxROP1^CA^-MxZR3.1. Identification of the strongest reaction to Fe deficiency responses in MxROP1^DN^-overexpressed plants indicates that MxROP1^DN^ plays a more important role in its interaction with MxZR3.1 during Fe deficiency responses. *In vivo* and *in vitro* experiments indicated that MxROP1 reduced the accumulation of MxZR3.1, and the MxZR3.1 protein was degraded by the 26S protein proteasome. Thus, it may be related to the regulation of ubiquitination degradation. The opposite roles of MxROP1^DN^ and MxZR3.1 in Fe deficiency responses and the degradation of MxZR3.1 protein by MxROP1 clearly showed that MxROP1 was induced to interact with MxZR3.1 to accelerate the degradation of its protein, thus, releasing its negative effects on Fe deficiency responses.

The classical transcriptional inhibitory domain EAR exists in the protein sequence of *MxZR3*.*1*, and this domain normally serves as an inhibitor in numerous pathways (Le et al., 2016). This is consistent with our results that TRV-*MxZR3*.*1* plants had enhanced responses to Fe deficiency. FIT and its activation complex bHLH39 regulate *FRO2* and *IRT1* transcription (Jeong et al., 2017). Here, we demonstrated that MxbHLH39 from *M. xiaojinensis* interacted with the MxZR3.1 protein. *bHLH039* is significantly upregulated under Fe deficiency, indicating that only a small amount of the bHLH039 protein is required when there is a sufficient supply of iron, which requires a mechanism to prevent the effects of excessive protein. This hypothesis is supported by the fact that the overexpression of bHLH039 when the amount of Fe is sufficient could overload the iron storage system and severely impact the accumulation of iron (Naranjo-Arcos et al., 2017). We hypothesize that MxZR3.1 plays a role in limiting the levels of bHLH39 proteins under a sufficient supply of iron. The addition of MxROP1 competed with bHLH39 to interact with ZR3.1. This suggests that under Fe deficiency conditions, a large amount of MxROP1^DN^ is induced to competitively bind to ZR3.1, which reduces the stability of ZR3.1 protein, thus, releasing the amount of bHLH39 that is bound. This should result in the regulation of the response to Fe deficiency. In addition, the increased transport activity of H^+^-ATPase and the expression of *MxHA2* in TRV-*MxZR3*.*1* plants under Fe deficiency indicates that *MxZR3*.*1* may negatively regulate the response to Fe deficiency by regulating *MxHA2* directly or by regulating transcription factors that are induced by Fe deficiency. However, further research is merited.

Based on our results in this study, we proposed a working model on MxROP1-MxZR3.1-MxbHLH39 to maintain Fe homeostasis in the apple rootstock *M. xiaojinensis* (Figure 7**F**). When plants are exposed to an environment deficient in iron, the non-activated state of MxROP1 is induced after sensing the Fe deficiency signal. MxROP1 in the GDP state complexes with the target protein MxZR3.1, so that MxZR3.1 is degraded through the 26S protein proteasome pathway. The bHLH39 protein is then released from MxZR3.1 to enhance the uptake of Fe. When the plants have sufficient iron, MxZR3.1 will be released from MxROP1^DN^ to bind to MxbHLH39, inhibiting the excess absorption of Fe.

## EXPERIMENTAL PROCEDURES

### Plant materials and growth conditions

The apple rootstock cultivars *M. xiaojinensis* Cheng et Jiang and *M. baccata* Borkh.) and the apple callus ‘Orin’ were used as the WT control to compare with diverse overexpression or RNAi plants in the relative background in the experiments. *M. xiaojinensis* seedlings were cultured for <u>p</u>hysiological and molecular analyses of the Fe-deprivation response based on one previous study (Sun et al., 2016). The surfaces of *M. baccata* seeds were washed with distilled water. The seeds were then placed in clean gauze and stratified for 30-40 day at 4 °C. The germinated seeds were planted in soil and vermiculite of 1:1 ratio for 7-10 days as materials to be injected. ‘Orin’ apple callus were cultured for <u>p</u>hysiological and molecular analyses of the Fe-deprivation response (Zhao et al., 2016).

For normal soil growth, tobacco seedlings were transferred to pots that contained a mixture of soil: roseite (1:1) and grown in the greenhouse under a 16-h/ 8-h light/dark and 25 °C/ 20 °C day/ night.

### Plasmid construction and generation of transgenic plants

All the primer pairs used are shown in Table S2. The seamless cloning Invitrogen (Carlsbad, CA, USA) was used to generate the vectors used in the Y2H, BiFC, pull-down, and LCI assays and plant transformations. The recombinant protein was expressed in plants using *A. tumefaciens* strain GV3101 mediated transformation (Xie et al., 2012), and *A. rhizogenes*-mediated transformation used strain K599 (Meng et al., 2019). The transgenic plants obtained were used for further analysis.

### Total RNA isolation and gene expression analysis

Total RNA was extracted using an improved CTAB method (Gasic et al., 2004) and treated with DNase I. cDNA was synthesized from 2 µg of total RNA using a TRUE Script Reverse Transcription Kit (+gDNA Eraser) (Invitrogen). Quantitative real-time PCR was performed on an ABI Quant Studio™ 6 Flex system (Applied Biosystems, Waltham, MA, USA) with SYBR Green PCR master mix (TaKaRa, Dalian, China). All the genes expressed were normalized against the internal control gene *MdACTIN* (CN938023). Three biological replicates were generated for each analysis. The primers used for real-time quantitative reverse transcription PCR (qRT-PCR) are shown in Table S2.

### Root ferric chelate reductase assay

The activity of FCR was determined as previously described (Grusak et al., 1995), and the amount of activity in callus was determined (Schikora and Schmidt, 2001). Briefly, the whole excised root and callus were placed in a tube filled with assay solution (0.5 mM CaSO4, 0.1 mM MES, 0.5 mM bathophenanthroline disulfonic acid disodium salt hydrate [BPDS] and 0.5 mM Fe-EDTA) for 1 h in the dark at room temperature. The absorbance of the assay solutions was measured at 535 nm by spectrophotometry (UV1800; Shimadzu, Kyoto, Japan), and the data were quantified using three biological replicates. The activity of FCR was visualized by placing roots and callus on culture plates that contained 1% agar and the same media as the assay solutions for 1 h in the dark.

### Rhizosphere acidification assay

Rhizosphere acidification was examined as previously described (Yi and Guerinot, 1996). Transgenic apple plants and callus were transferred to a 1% agar plate that contained 0.006% bromocresol purple and 0.2 mM CaSO4 for 24 h at a pH of 6.5 that had been adjusted with 1 M NaOH.

### Perls blue staining

To localize Fe^3+^, the leaves and roots of WT and transgenic apple lines were incubated with Perls stain solution (equal volumes of 4% [v/v] HCl and 4% [w/v] potassium ferrocyanide) for 60 min.

### Y2H

Y2H protein interaction assays were performed according to the manufacturer’s instructions as described in the Yeast Protocols Handbook (Clontech; TaKaRa Bio USA, Inc., Forster, CA, USA). Constructs that contained NZR3.1, CZR3.1, C1ZR3.1, C2ZR3.1 and ZR3.1 were fused to GAL4 AD and ROPs, ROP1^CA^, and ROP1^DN^, bHLH39-1 and bHLH39-2 fused to the BD were co-transformed into yeast Y2HGold (Clontech). Transformants were grown on SD-Leu-Trp plates, and then picked as primary positives and transferred to SD-Leu-Trp-His selection plates for protein interaction studies.

### BiFC assay

The vectors that were used in the BiFC assays to form different combinations of constructs were transformed into *A. tumefaciens* strain GV3101 and then co-injected into young leaves of *Nicotiana benthamiana*. After 48 h, the fluorescence was observed by a Nikon D-ECLIPSE C1 spectral confocal laser-scanning system (Tokyo, Japan).

### Pull-Down Analysis

These constructs were expressed in *Escherichia coli* BL21 (DE3), and the fusion proteins were purified using the corresponding agarose beads. GST, GST-ROP1 or GST-bHLH39s was eluted from the glutathione-agarose beads and then incubated with His-ZR3.1 for 4 h at 4°C, washed thoroughly and boiled in 5× SDS-PAGE sample buffer for 10 min.

### LCI assays

The cLUC-ROP1, cLUC-ROP1^CA^, cLUC-ROP1^DN^ and nLUC-ZR3.1 constructs were generated as shown in Table S1. Agrobacteria that harbored different combinations of constructs were co-infiltrated into *N. benthamiana*, and the infiltrated leaves were analyzed for LUC activity using LUCK2019 imaging (LB985) after 48 h of infiltration (Zhao et al., 2020).

### Protein extraction and western blotting

A total of 2 g of transgenic apple callus or apple plants for each sample were ground in a protein extraction reagent buffer (Cwbio, Jiansu, China) that contained a protease inhibitor cocktail. Protein extracts were boiled in 5× SDS-PAGE sample buffer and analyzed by immunoblotting using anti-His and anti-GST antibodies.

### Cell-free degradation

His-MxZR3.1 protein was induced and purified *in vitro*. The total proteins of the *35S::ROP1-GFP* callus were subsequently extracted in degradation buffer (25 mM Tris-HCl, pH 7.5, 10 mM NaCl, 10 mM MgCl_2_, 4 mM phenylmethylsulfonyl fluoride [PMSF], 5 mM DTT, and 10 mM ATP) as described by Wang et al (2009). Each reaction mix contained 100 ng of His-MdbHLH104 and 500 mg of total protein from *35S::ROP1-GFP* callus. A total of 50 mM MG132 was added 30 min before the proteasome inhibitor experiments. The reaction mixes were incubated at 22°C, terminated at different times and then boiled for 10 min.

### Statistical analysis

The data were analyzed using GraphPad Prism 8 (San Diego, CA, USA) using a one-way analysis of variance (ANOVA) (Duncan correction) or *t*-test (P < 0.05), and statistically significant differences were indicated by different lowercase letters or *, respectively.

## Data availability

The data supporting the findings of this study are available from the corresponding author, Yi Wang, upon request.

## Acknowledgements

This work was supported by the National Natural Science Foundation of China (32172537 and 31972385), the earmarked fund for China Agriculture Research System (CARS-27), the Construction of Beijing Science and Technology Innovation and Service Capacity in Top Subjects (CEFF-PXM2019_014207_000032), the 2115 Talent Development Program of China Agricultural University, and the Key Laboratory of Beijing Municipality of Stress Physiology and Molecular Biology for Fruit Trees. We thank Professors Yujin Hao of Shandong Agricultural University and Shuhua Yang of China Agricultural University for providing vectors and technical support throughout this study.

## Author contributions

KL and LZ wrote the manuscript; KL performed most of the experiments; LJ and LZ helped to clone genes, express proteins and vector constructions; QS helped to perform the pulldown assays; YF, TW, XX and XZ gave advice during the data analysis and helped with the phenotypic analysis. YW and ZH conceived the project and edited the manuscript.

## SUPPORTING INFORMATION

**Fig. S1** Identification of MxROP1.

**Fig. S2** Identification of *MxROP1, MxROP1*^*DN*^ and *MxROP1*^*CA*^ transgenic apple roots.

**Fig. S3** Phenotypic and GFP signal analysis of *MxROP1, MxROP1*^*DN*^ and *MxROP1*^*CA*^ transgenic apple roots.

**Fig. S4** Y2H assay showed that MxROP1 interacts with candidate target protein.

**Fig. S5** Y2H assay between MdZRs and MxROPs.

**Fig. S6** Iron deficiency induces the expression of *MxZR3*.*1*.

**Fig. S7** Relative expression of Fe deficiency genes in silencing *MxZR3*.*1* lines under +Fe or -Fe.

**Fig. S8** Relative expression of Fe deficiency genes in transgenic callus under +Fe or -Fe.

**Table S1** Physical and chemical characters of MdROPs family members.

**Table S2** Primers used in this study.

